# Visual Interpretation of the Meaning of *k*_cat_/*K*_M_ in Enzyme Kinetics

**DOI:** 10.1101/2021.12.09.471995

**Authors:** Chiwook Park

## Abstract

*k*_cat_ and *k*_cat_/*K*_M_ are the two fundamental kinetic parameters in enzyme kinetics. *k*_cat_ is the first-order rate constant that determines the reaction rate when the enzyme is fully occupied at a saturating concentration of the substrate. *k*_cat_/*K*_M_ is the second-order rate constant that determines the reaction rate when the enzyme is mostly free at a very low concentration of the substrate. Both parameters provide critical information on how the enzyme lowers the energy barriers along the reaction pathway for catalysis. However, it is surprising how often *k*_cat_/*K*_M_ is used inappropriately as a composite parameter derived by dividing *k*_cat_ with *K*_M_ to assess both catalytic power and affinity to the substrate of the enzyme. The main challenge in explaining the true meaning of *k*_cat_/*K*_M_ is the difficulty to demonstrate how the reaction energetics of enzyme catalysis determines *k*_cat_/*K*_M_ in a simple way. Here, I report a step-by-step demonstration on how to visualize the meaning of *k*_cat_/*K*_M_ on the reaction energy diagram. By using the reciprocal form of the expression of *k*_cat_/*K*_M_ with the elementary rate constants in kinetic models, I show that *k*_cat_/*K*_M_ is a harmonic sum of several kinetic terms that correspond to the heights of the transition states relative to the free enzyme. Then, I demonstrate that the height of the highest transition state has the dominant influence on *k*_cat_/*K*_M_, i. e. the step with the highest transition state is the limiting step for *k*_cat_/*K*_M_. The visualization of the meaning of *k*_cat_/*K*_M_ on the reaction energy diagram offers an intuitive way to understand all the known properties of *k*_cat_/*K*_M_, including the Haldane relationship.

## Introduction

When an enzyme catalyzes the conversion of a substrate to a product, the dependence of the reaction rate (*v*) on the substrate concentration ([S]) follows the following simple rate law:^1^

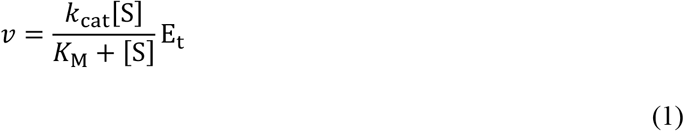

which is known as the Michaelis–Menten equation. When the substrate concentration is much greater than *K*_M_, i.e. [S] >> *K*_M_, Eq. 1 is simplified as

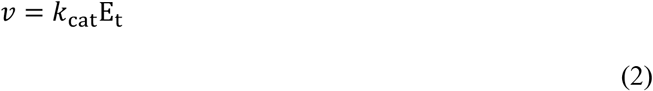

Under this condition, the reaction is first-order; the reaction rate is dependent only on the total enzyme concentration, not on the substrate concentration (Figure 1). Therefore, *k*_cat_ is a first-order rate constant with the unit of s^-1^ and frequently called the ‘turnover number’, the number of substrate molecules that one enzyme can catalyze per unit time. Also, *k*_cat_E_t_ is the maximum reaction rate (*V*_max_) that the enzyme can achieve under the given experimental condition. When the substrate concentration is much less than *K*_M_, i.e. [S] << *K*_M_, Eq. 1 is simplified as

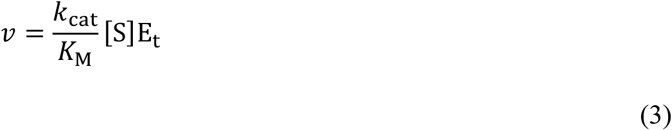

**Figure 1.**
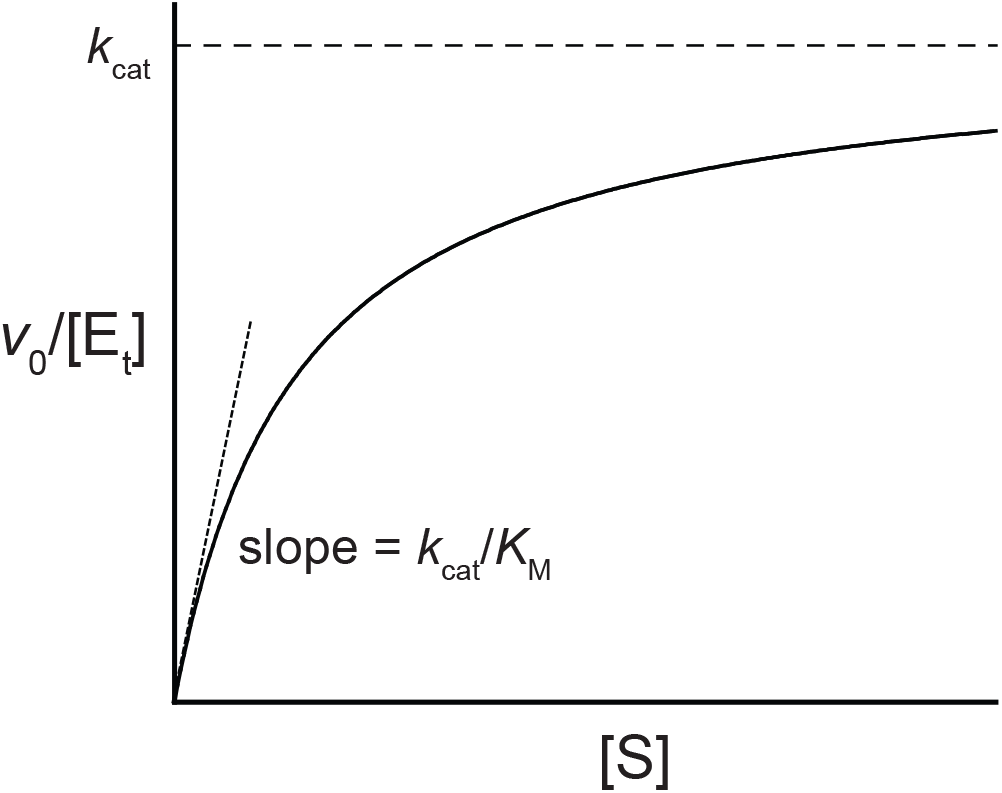
Plot of *v*/E_t_ versus [S] according to the Michaelis–Menten equation. At saturating concentrations of [S], *v*/E_t_ approaches at *k*_cat_. At low concentrations of [S], *v*/E_t_ increases linearly with the slope of *k*_cat_/*K*_M_ as [S] increases.

Under this condition, the reaction is second-order; the reaction rate is proportional to both substrate concentration and enzyme concentration (Figure 1). Therefore, *k*_cat_/*K*_M_ is a second-order kinetic constant with the unit of M^-1^s^-1^. *k*_cat_/*K*_M_ is frequently called the ‘specificity number’ because *k*_cat_/*K*_M_ varies greatly with different substrates. The Michaelis constant (*K*_M_) is just the ratio of *k*_cat_ to *k*_cat_/*K*_M_ and corresponds to the concentration of the substrate at which half of the enzyme is saturated. Enzymologists commonly use the two fundamental kinetic parameters, *k*_cat_ and *k*_cat_/*K*_M_, to investigate the mechanism of enzymatic catalysis (1, 5, 6). Most of all, *k*_cat_ and *k*_cat_/*K*_M_ allow them to prove the nature of the rate-limiting step in the enzyme-catalyzed reaction. Frequently, the changes in *k*_cat_ and *k*_cat_/*K*_M_ are monitored in varying assay conditions (pH, temperature, salts, viscosity, or the presence of an inhibitor or an activator), with different substrates, or with variants of enzymes to dissect the detailed mechanism of the enzymatic catalysis. Being the ratio of the two fundamental kinetic constants, the effects of the change in the experimental condition on *K*_M_ is quite complex and usually do not provide any interpretable or meaningful information on enzymatic catalysis (1, 5).

While it has been well accepted among enzymologists that *k*_cat_ and *k*_cat_/*K*_M_ are the two fundamental kinetic parameters for enzymatic catalysis, a general misunderstanding of *k*_cat_/*K*_M_ is still prevalent in literature and even in biochemistry textbooks. Frequently, *k*_cat_ and *K*_M_ are considered as the two primary kinetic parameters one can obtain from the Michaelis-Menten equation, and *k*_cat_/*K*_M_ is considered as the ratio of *k*_cat_ to *K*_M_, which is mathematically true, but tremendously misleading. Considering *k*_cat_/*K*_M_ as a composite parameter greatly hinders the understanding of the true meaning of *k*_cat_/*K*_M_ and the proper interpretation of *k*_cat_/*K*_M_ in studying enzymatic catalysis. The reason for the confusion of *k*_cat_/*K*_M_ as a composite parameter is partly because the symbol of the kinetic parameter has the form of the ratio of *k*_cat_ to *K*_M_. To avoid this issue, a new symbol *k*_cap_ (capture constant) has been proposed to replace *k*_cat_/*K*_M_ (5), but unfortunately, the new notation has not been widely accepted. However, the major challenge regarding *k*_cat_/*K*_M_ is the difficulty to figure out the relationship between *k*_cat_/*K*_M_ and the elementary rate constants from the expression of *k*_cat_/*K*_M_ (see Eq. 8 below as an example). It has been shown from the mathematical expression that *k*_cat_/*K*_M_ is the ‘net’ rate constant for the substrate association step (E + S → ES) (3) or the apparent rate constant for the ‘capture’ of the substrate (5), reporting the kinetics of the formation of the enzyme-substrate complex (ES) that eventually “goes on to produce the product” (1) or “are destined to yield the product” (7). However, to a non-expert in enzyme kinetics, these explanations are not so easy to understand either.

Here I report a simpler way to explain the meaning of *k*_cat_/*K*_M_ visually on the reaction energy diagram using the reciprocal expressions of *k*_cat_/*K*_M_. First, using the reciprocal expressions of *k*_cat_/*K*_M_, I show that *k*_cat_/*K*_M_ is a harmonic sum of several kinetic terms, i.e. the reciprocal form of *k*_cat_/*K*_M_ is the sum of the reciprocal forms of a group of kinetic terms. This approach to express *k*_cat_/*K*_M_ as a harmonic sum of several kinetic terms has been used in literature (8, 9), but its value in deciphering the meaning of *k*_cat_/*K*_M_ has not been fully explored. Next, I examine the meaning of each kinetic term comprising *k*_cat_/*K*_M_ using the reaction energy diagram and explain that each term actually corresponds to the height of the barrier corresponding to an individual step in the kinetic model. This analysis clearly shows that *k*_cat_/*K*_M_ is determined by the heights of the transition states relative to E + S and influenced (or limited) dominantly by the highest transition state. Using this approach, I also demonstrate that the previously known properties of *k*_cat_/*K*_M_, including the Haldane relationship, can be explained based on reaction energetics.

### Reciprocal Expression for *k*_cat_/*K*_M_

Here is a minimal, but realistic kinetic model for an enzyme-catalyzed reaction:^2^

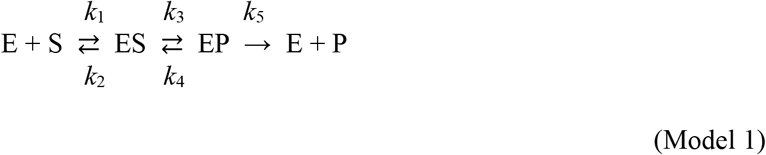

For simplicity, I assume that the reaction has only one substrate and one product. This model contains five rate constants. While *k*_2_, *k*_3_, *k*_4_, and *k*_5_ are first-order rate constants, *k*_1_ is a second-order rate constant for the substrate association. The product-release step is reversible in many enzyme-catalyzed reactions; however, we usually make the kinetic studies simpler by setting up the reaction only with the substrate and the enzyme (without the product), i.e. [P] = 0. Under this condition, the reverse reaction (P → S) is negligible, and only the forward reaction (S → P) determines the overall reaction rate, which allows us to determine *k*_cat_ and *k*_cat_/*K*_M_ for the forward reaction without any unnecessary complexity. Moreover, the inclusion of the irreversible step makes the derivation of the rate equation much easier. I will revisit later the case of enzymes that work under the condition that the substrate and the product are in equilibrium.

By using the steady-state approximation, we derive the rate equation for Model 1 as:^3^

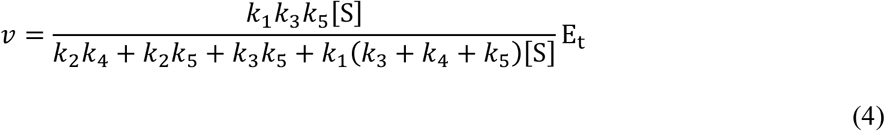

When the substrate concentration is much greater than *K*_M_, i.e. [S] >> *K*_M_, the rate is determined by *k*_cat_ (Eq. 2) and Eq. 4 is simplified as:

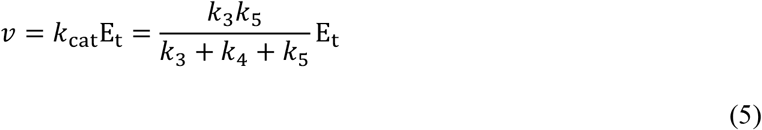

Therefore,

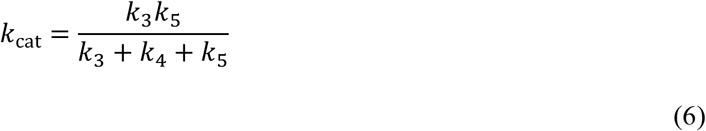

When the substrate concentration is much less than *K*_M_, i.e. [S] << *K*_M_, the rate is determined by *k*_cat_/*K*_M_ (Eq. 3), and Eq. 4 is simplified as

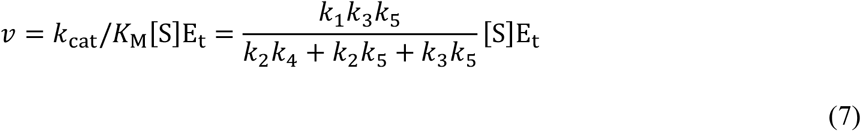

Therefore,

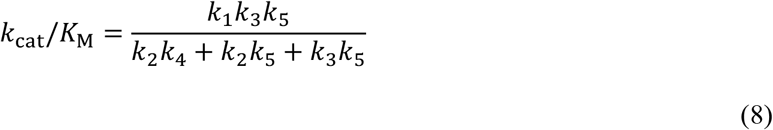

Even with this minimal kinetic model, it is difficult to figure out directly from Eq. 8 how each step in the kinetic model contributes to *k*_cat_/*K*_M_. The difficulty to decipher the mathematical expression of *k*_cat_/*K*_M_ like Eq. 8 is a potential factor causing the prevalent misconception of *k*_cat_/*K*_M_ as a composite kinetic parameter (a ratio of *k*_cat_ to *K*_M_) rather than a fundamental kinetic parameter.^4^

The reciprocal form of Eq. 8 offers us a much simpler way to demonstrate the meaning of *k*_cat_/*K*_M_.

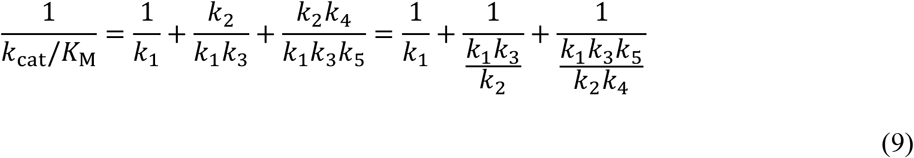

According to Eq. 9, *k*_cat_/*K*_M_ is actually the harmonic sum (the reciprocal of the sum of the reciprocals of each member) of three kinetic terms: *k*_1_, *k*_1_*k*_3_/*k*_2_, and *k*_1_*k*_3_*k*_5_/*k*_2_*k*_4_. This expression shows that *k*_cat_/*K*_M_ is ‘mathematically’ analogous to the total resistance of a parallel circuit composed of resistance of *k*_1_, *k*_1_*k*_3_/*k*_2_, and *k*_1_*k*_3_*k*_5_/*k*_2_*k*_4_. Note that in reality the rate constant is opposite to the resistance; the greater the rate constant is, the faster the reaction is. According to the mathematical relationship shown in Eq. 9, the smaller value of *k*_1_, *k*_1_*k*_3_/*k*_2_, and *k*_1_*k*_3_*k*_5_/*k*_2_*k*_4_ has the greater influence on *k*_cat_/*K*_M_. Moreover, *k*_cat_/*K*_M_ is always smaller than the smallest of the three terms and converges to the smallest term when the other two terms are significantly greater than the smallest term (Table 1). It is analogous to the fact that in a parallel electric circuit the current flows most through the path with the smallest resistance, which has the dominant influence on the overall resistance.

**Table 1.** Influence of individual kinetic terms on *k*_cat_/*K*_M_ in Eq. 9. *k*_1_ is fixed at 10^6^ M^-1^s^-1^, and *k*_1_*k*_3_/*k*_2_ and *k*_1_*k*_3_*k*_5_/*k*_2_*k*_4_ are varied from 10^5^ M^-1^s^-1^ to 10^7^ M^-1^s^-1^. The least terms, which have the dominant influence on *k*_cat_/*K*_M_, are shown in bold.

| $k_1$<br>($\times 10^6 \text{ M}^{-1}\text{s}^{-1}$) | $k_1k_3/k_2$<br>($\times 10^6 \text{ M}^{-1}\text{s}^{-1}$) | $k_1k_3k_5/k_2k_4$<br>($\times 10^6 \text{ M}^{-1}\text{s}^{-1}$) | $k_{\text{cat}}/K_{\text{M}}$<br>($\times 10^6 \text{ M}^{-1}\text{s}^{-1}$) |
| --- | --- | --- | --- |
| 1 | **0.1** | **0.1** | 0.048 |
| 1 | **0.1** | 1 | 0.083 |
| 1 | **0.1** | 10 | 0.09 |
| 1 | 1 | **0.1** | 0.083 |
| **1** | **1** | **1** | 0.33 |
| **1** | **1** | 10 | 0.48 |
| 1 | 10 | **0.1** | 0.09 |
| **1** | 10 | **1** | 0.48 |
| **1** | 10 | 10 | 0.83 |

Now let’s figure out the meanings of *k*_1_, *k*_1_*k*_3_/*k*_2_, and *k*_1_*k*_3_*k*_5_/*k*_2_*k*_4_. The meaning of *k*_1_ is obvious; by definition, *k*_1_ is the rate constant for the formation of the ES complex. Deciphering the meanings of *k*_1_*k*_3_/*k*_2_ and *k*_1_*k*_3_*k*_5_/*k*_2_*k*_4_ requires the consideration of the reaction energetics. The reaction energy diagram (Figure 2) shows the relative height of the barrier for each step in Model 1. For a first-order rate constant, the corresponding barrier is obviously Δ*G*‡ of the transition-state theory, which is linearly proportional to ln(1/k); thus, the smaller the rate constant, the higher the barrier is. To include *k*_1_, a second-order rate constant, in the reaction energy diagram with first-order rate constants together, one needs to convert *k*_1_ to a pseudo-first-order rate constant by multiplying by [S].^5^ Therefore, the first barrier from E+S in the reaction energy diagram is determined by *k*_1_[S]. We make the level of E + S much lower than ES or EP in the reaction energy diagrams because *k*_cat_/*K*_M_ determines the kinetics when the substrate concentration is low, i.e. most of the enzyme is in the free from (E). In the reaction energy diagram, *k*_1_, *k*_1_*k*_3_/*k*_2_ and *k*_1_*k*_3_*k*_5_/*k*_2_*k*_4_ have clear physical meanings. As shown in Figure 2, *k*_1_, *k*_1_*k*_3_/*k*_2_ and *k*_1_*k*_3_*k*_5_/*k*_2_*k*_4_, when multiplied by [S], correspond to the heights of the transition states relative to E + S for the substrate association step (E + S → ES), the chemical conversion step (ES → EP), and the product release step (EP → E+P), respectively.^6^ Therefore, the reciprocal expression of *k*_cat_/*K*_M_ (Eq. 9) shows that *k*_cat_/*K*_M_ is determined by the heights of the three transition states relative to E + S.

**Figure 2.**
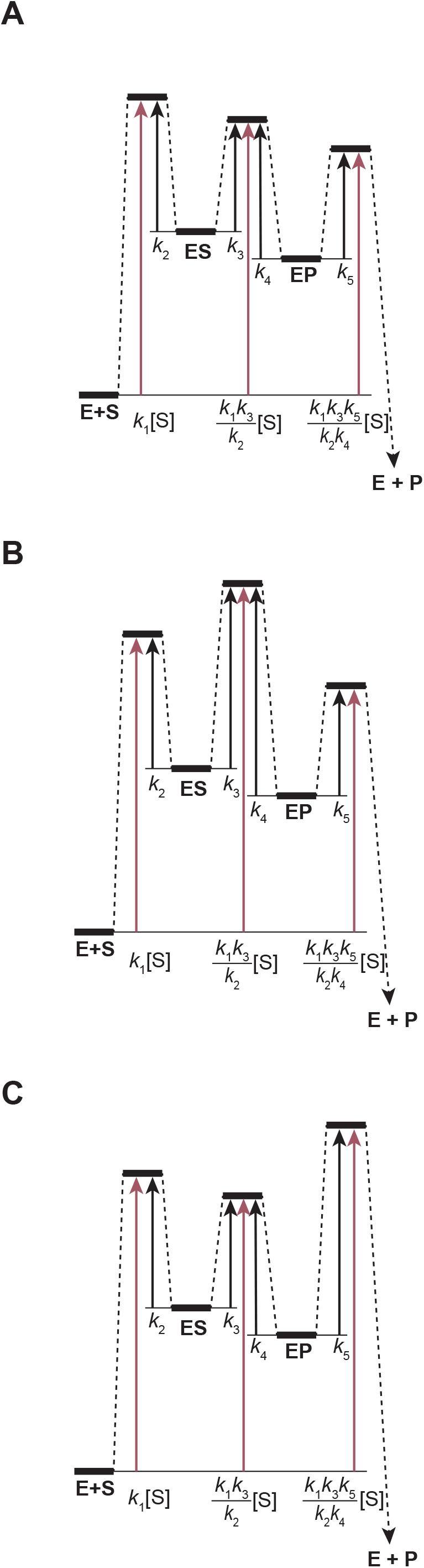
Normative reaction energy diagrams for the kinetic model shown in Model 1. The heights of the transition states corresponding to *k*_1_, *k*_1_*k*_3_/*k*_2_, and *k*_1_*k*_3_*k*_5_/*k*_2_*k*_4_ relative to E + S are indicated by teal arrows. **(A)** The transition state for the substrate-association step (E + S → ES) is the highest. **(B)** The transition state for the chemical-conversion step (ES → EP) is the highest. **(C)** The transition state for the product-release step (EP → E + P) is the highest.

### Rate-limiting step for *k*_cat_/*K*_M_

Figure 2 shows three versions of the reaction energy diagrams for Model 1. The substrate association step (E + S → ES), the chemical conversion step (ES → EP), and the product release step (EP → E+P) has the highest transition state in Figures 2A, 2B, and 2C, respectively. Because the height of the transition state is linearly proportional to ln(1/k), the least of *k*_1_, *k*_1_*k*_3_/*k*_2_ and *k*_1_*k*_3_*k*_5_/*k*_2_*k*_4_ has the highest transition state and also the greatest influence on *k*_cat_/*K*_M_. In other words, the step with the highest transition state in each reaction energy diagram in Figure 2 is the rate-limiting step for *k*_cat_/*K*_M_.^7^ In reverse, measuring of *k*_cat_/*K*_M_ of an enzyme-catalyzed reaction is a way to probe the highest transition state of the reaction. Note that the relative heights of the transition states do not change when the substrate concentration is varied. When the substrate concentration is increased, only the level of E + S on the reaction energy diagrams moves up, and the relative heights of the transition states remain same. Therefore, information obtained from *k*_cat_/*K*_M_ on the relative heights of the transition states is still valid at any substrate concentration.^8^

When the transition state for the substrate association step (E + S → ES) is the highest (Figure 2A), *k*_cat_/*K*_M_ is determined mostly by the kinetics of the substrate association (*k*_cat_/*K*_M_ ~ *k*_1_). When the enzyme catalyzes the chemical conversion efficiently, the substrate association step frequently becomes the rate-limiting step for *k*_cat_/*K*_M_. Because the substrate association cannot be faster than the collision between the enzyme and the substrate by diffusion, the kinetics of diffusion is the ultimate physical limit to *k*_cat_/*K*_M_. When *k*_cat_/*K*_M_ reaches to this physical limit, *k*_cat_/*K*_M_ is said to be diffusion-controlled (10). Depending on the size of the substrate, *k*_cat_/*K*_M_ can reach up to 10^8^–10^9^ M^-1^s^-1^. Some enzymes, such as fumarase (10) and triose phosphate isomerase (11), are known to have *k*_cat_/*K*_M_ values that approach this limit. The substrate association is normally much slower than diffusion because it is not likely that every collision leads to the formation of the enzyme–substrate complex capable for catalysis (ES) (12, 13).

When the transition state for the chemical conversion (ES → EP) or the product release (EP → E + P) is the highest (Figures 2BC), *k*_cat_/*K*_M_ is determined mostly by *k*_1_*k*_3_/*k*_2_ or *k*_1_*k*_3_*k*_5_/*k*_2_*k*_4_, respectively. As *k*_2_/*k*_1_ is the dissociation equilibrium constant for the ES complex (*K*_S_), *k*_cat_/*K*_M_ in Figure 2B is approximated as *k*_3_/*K*_S_. In this case, *k*_cat_/*K*_M_ is indeed the measure of both catalytic power (*k*_3_) and affinity to the substrate (*K*_S_). Even though this relationship is valid only when the transition state for the chemical conversion is the highest, it is so commonly used as the general meaning of *k*_cat_/*K*_M_. Because enzymes in nature are highly evolved to catalyze the chemical conversion, it is somewhat rare that the transition state for the chemical conversion is the highest with their physiological substrates. Therefore, the interpretation of *k*_cat_/*K*_M_ as the measure of both catalytic power (*k*_3_) and affinity to the substrate (*K*_S_) should be reserved for poor substrates or catalytically impaired variants of enzymes. Also, as commonly as *k*_cat_/*K*_M_ is limited by the substrate association, *k*_cat_/*K*_M_ can be limited by the product release (see the discussion on the Haldane relationship below).

### Generalized kinetic model

Now we derive the reciprocal expression of *k*_cat_/*K*_M_ for a generalized kinetic model even without deriving the rate equation. Let’s say the generalized kinetic model contains the free enzyme (E), *n* bound forms of the enzyme (E_b_’s), one substrate (S), and one product (P):

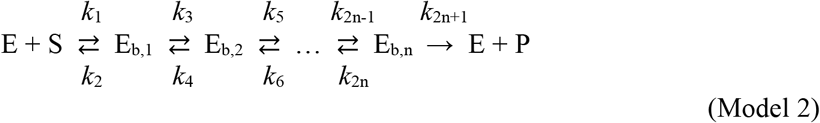

The kinetic model ends with an irreversible step, which leads to E + P. The reciprocal form of *k*_cat_/*K*_M_ of this generalized kinetic model is expressed as the sum of the reciprocals of rate constants corresponding to each transition state:

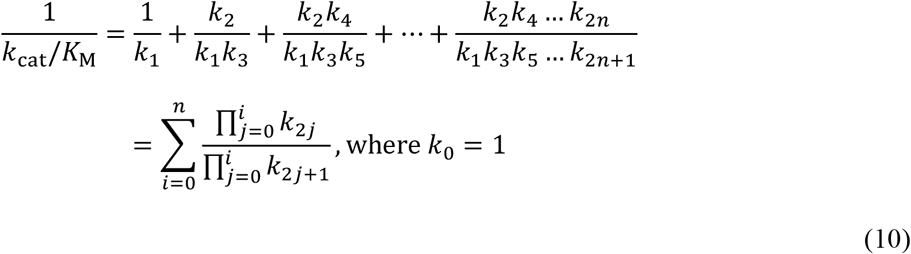

This generalized description of *k*_cat_/*K*_M_ clearly demonstrates that, in any kinetic model, *k*_cat_/*K*_M_ has a straightforward physical meaning, which can be easily shown on the reaction energy diagram. As stated above, the highest transition state out of the *n*+1 transition states in the reaction energy diagram has the dominant influence on *k*_cat_/*K*_M_.

### Steps that are considered in *k*_cat_/*K*_M_

It is commonly stated that the expression of *k*_cat_/*K*_M_ includes all the elementary rate constants from the free enzyme (E + S) to the first irreversible step (4). It is quite simple to demonstrate this property using the reciprocal expression of *k*_cat_/*K*_M_. By converting the reversible chemical conversion step (ES ⇄ EP) in Model 1 to an irreversible one (ES → EP), we can make Model 3:

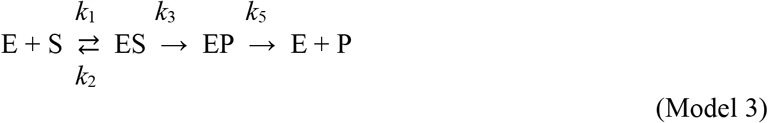

in which we do not have *k*_4_, which is the rate constant for the reverse reaction of the chemical conversion step (EP → ES). Eliminating *k*_4_ in Model 2 is mathematically identical to setting *k*_4_ to 0 in the expression of *k*_cat_/*K*_M_ for Model 2 (Eq. 9). Therefore, the reciprocal form of *k*_cat_/*K*_M_ for Model 3 is:

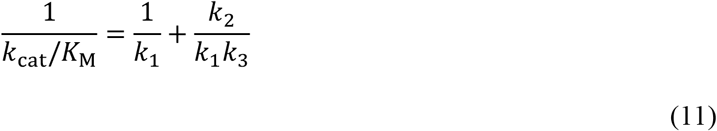

As we stated above, *k*_cat_/*K*_M_ in Eq. 11 includes the elementary rate constants from the free enzyme to the first irreversible step (EP → ES) in Model 3. We can also demonstrate the property with the generalized model (Model 2). In the reciprocal form of *k*_cat_/*K*_M_ for the generalized model (Eq. 10), all the rate constants for the reverse reactions (rate constants with even subscripts, such as *k*_2_, *k*_4_, *k*_6_, etc.) are in the numerators of the terms. By setting any of the rate constants with an even subscript to 0, the first term that contains the rate constant and all the terms following disappear from the expression of *k*_cat_/*K*_M_.

Any steps after the first irreversible step does not affect *k*_cat_/*K*_M_ because *k*_cat_/*K*_M_ describes the kinetics from E + S, and any species after the first irreversible step cannot go back to E + S. We can explain this more intuitively using the reaction energy diagram. The heights of the transition states determine *k*_cat_/*K*_M_. An irreversible step is where the rate constant for the reverse reaction is 0. Therefore, an irreversible step has an infinite drop in the forward direction in the reaction energy diagram, which we indicate as a downward arrow (Figure 2). Due to this infinite drop, transition states after the first irreversible step are infinitely lower than those before the irreversible step and cannot make any contribution to *k*_cat_/*K*_M_.

### Barriers versus Troughs

Reaction energy diagrams for enzyme-catalyzed reactions are composed of barriers (transition states) and troughs (bound forms, such as ES and EP). It is known that the depths of the troughs do not affect *k*_cat_/*K*_M_ (4). We already demonstrated above that *k*_cat_/*K*_M_ is determined by the heights of the transition states. When the depth of any trough varies without affecting the heights of the neighboring barriers (Figure 3A), *k*_cat_/*K*_M_ stays the same.^9^ In other words, *k*_cat_/*K*_M_ reports how effectively the enzyme stabilizes the transition states, regardless of its stabilization of the bound forms (or affinities to the substrate, the product, or any intermediates in between). One can verify this using the reciprocal expression of *k*_cat_/*K*_M_. When the energy level of ES is raised (Figure 3A), both *k*_2_ and *k*_3_ are increased to the same degree, which means that the ratio of *k*_2_ to *k*_3_ (*k*_2_/*k*_3_) remains unaffected. As one can verify in Eqs. 9 and 11, *k*_cat_/*K*_M_ does not change when *k*_2_/*k*_3_ remains constant.

**Figure 3.**
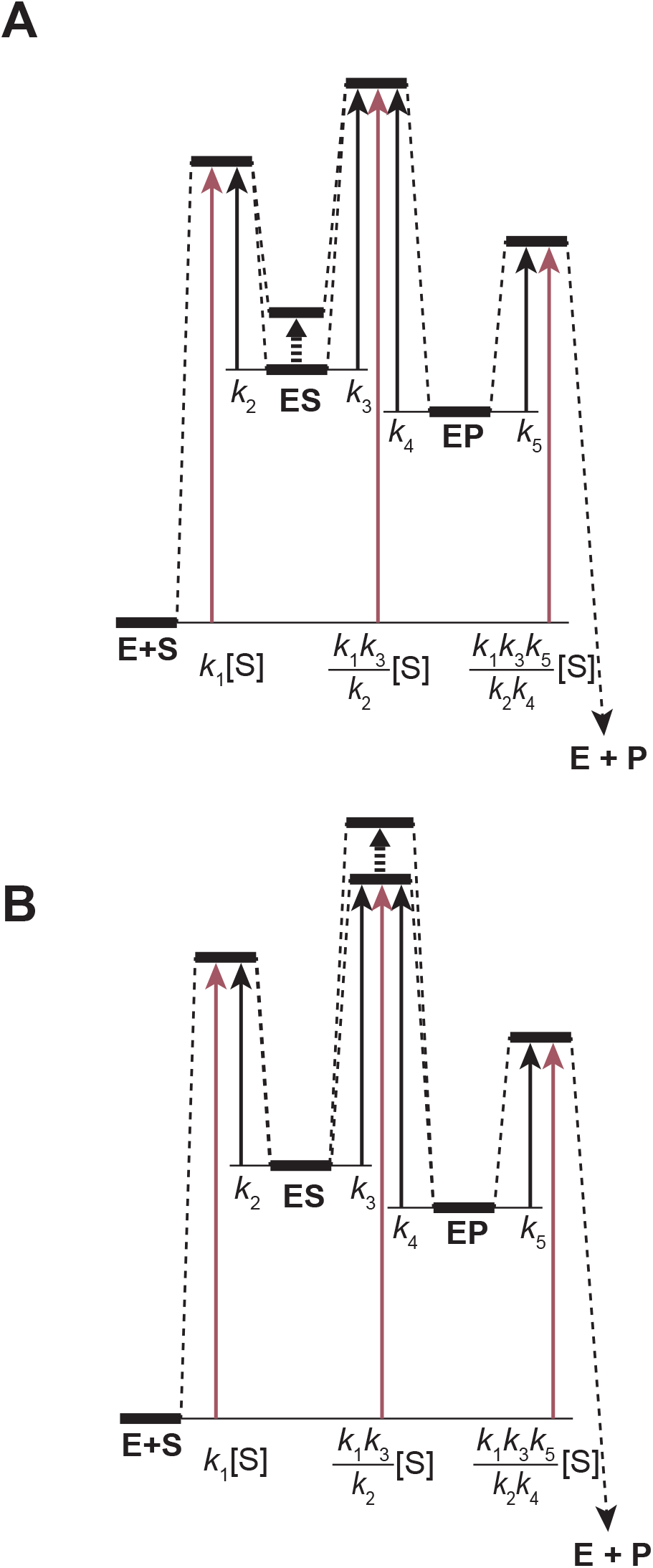
Effects of the change in the energy levels of a trough and a barrier on *k*_cat_/*K*_M_. The heights of the transition states corresponding to *k*_1_, *k*_1_*k*_3_/*k*_2_, and *k*_1_*k*_3_*k*_5_/*k*_2_*k*_4_ relative to E + S are indicated by teal arrows. **(A)** The energy level of the ES complex is increased. **(B)** The energy level of the transitions state for the chemical conversion step (ES → EP) is increased.

Now we consider the case that the height of only one of the barriers is increased. Such a case is common in kinetic studies on isotope effects (1). When a covalent bond breaks or forms during the chemical conversion step (ES ⇄ EP), the replacement of an atom with a heavier isotope selectively slows the chemical conversion step (Figure 3B). In that case, the height of the transition state is increased, and *k*_cat_/*K*_M_ may decrease. However, the degree of the change in *k*_cat_/*K*_M_ depends on the degree of the influence of the chemical conversion step on *k*_cat_/*K*_M_. If the transition state of the chemical conversion step is the highest barrier, the isotope effect on *k*_cat_/*K*_M_ is apparent (Figure 3B). If the transition state of the chemical conversion step is not the highest barrier, the isotope effect on *k*_cat_/*K*_M_ is significantly suppressed or even undetectable because *k*_cat_/*K*_M_ is determined mostly by a transition state that is not affected by the presence of the isotope.

### Haldane Relationship

Many of the enzyme-catalyzed reactions are reversible, and enzymes can catalyze both forward and reverse reactions. Moreover, the physiological function of some enzymes, such as adenylate kinase and triose phosphate isomerase, is to maintain the equilibrium between the substrate and the product in cells. For reversible reactions catalyzed by an enzyme, the following relationship holds true:

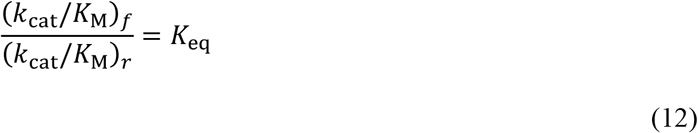

where (*k*_cat_/*K*_M_)_f_ is *k*_cat_/*K*_M_ for the forward reaction, (*k*_cat_/*K*_M_)_r_ is *k*_cat_/*K*_M_ for the reverse reaction, and *K*_eq_ is the equilibrium constant of the reaction. This relationship was originally discovered by Haldane and known to be the Haldane relationship (14). As long as only a catalytic amount of an enzyme is present, the presence of an enzyme does not affect *K*_eq_. Eq. 12 shows that *k*_cat_/*K*_M_ for the forward reaction and *k*_cat_/*K*_M_ for the reverse reaction are mathematically tied together by the equilibrium constant. When one of the values is increased or decreased, the other also varies in the same direction proportionally. Therefore, the Haldane relationship is another constraint on the limit of *k*_cat_/*K*_M_. For example, if *K*_eq_ is significantly smaller than 1, *k*_cat_/*K*_M_ for the forward reaction cannot reach to the diffusion limit. According to Eq. 12, when *K*_eq_ is significantly smaller than 1, (*k*_cat_/*K*_M_)_r_ is much greater than (*k*_cat_/*K*_M_)_f_. If (*k*_cat_/*K*_M_)_f_ is diffusion-controlled, (*k*_cat_/*K*_M_)_r_ should be greater than the diffusion limit, which is physically impossible. Likewise, if *K*_eq_ is significantly larger than 1, *k*_cat_/*K*_M_ for the reverse reaction cannot reach the diffusion limit.

The Haldane relationship can be understood more intuitively by using the reaction energy diagram. Let’s construct the kinetic model for a reversible reaction by adding the rate constant for the product association (*k*_6_) to Model 1:

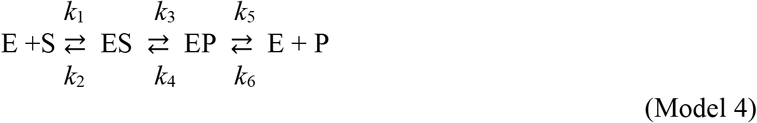

To experimentally determine (*k*_cat_/*K*_M_)_f_ for a reversible reaction, one keeps the product concentration close to 0 so that the reverse reaction is negligible. This experimental condition is mathematically identical to setting *k*_6_ to 0, which makes the kinetic model same as Model 1 and the reciprocal form of (*k*_cat_/*K*_M_)_f_ same as Eq. 9:

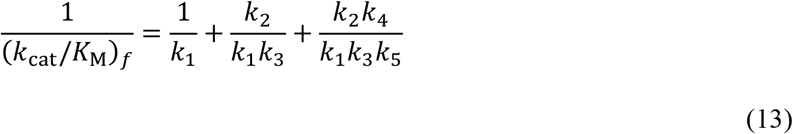

Likewise, we determine the expression for (*k*_cat_/*K*_M_)_r_ by setting *k*_1_ to 0:

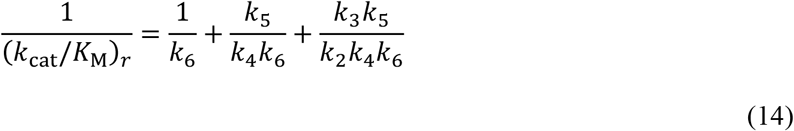

Now let’s assume that one transition state is significantly higher than others. While Haldane relationship does not require this assumption, we can use this assumption to simplify the demonstration. It is obvious that the highest transition state for the forward reaction is also the highest transition state for the reverse reaction. The same step has the dominance influence on both (*k*_cat_/*K*_M_)_f_ and (*k*_cat_/*K*_M_)_r_. In other words, the same step is rate-limiting for both (*k*_cat_/*K*_M_)_f_ and (*k*_cat_/*K*_M_)_r_. In case of the reaction energy diagram in Figure 4, the chemical conversion step (ES ⇄ EP) has the highest transition state, and (*k*_cat_/*K*_M_)_f_ and (*k*_cat_/*K*_M_)_r_ can be approximated as:

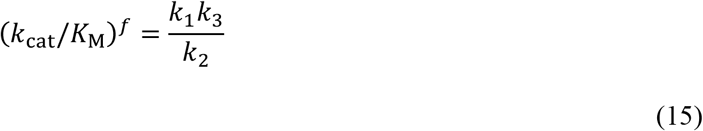

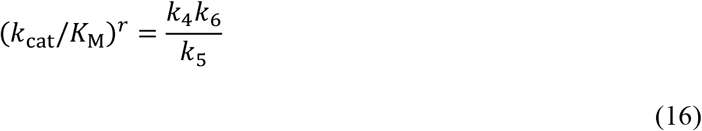

**Figure 4.**
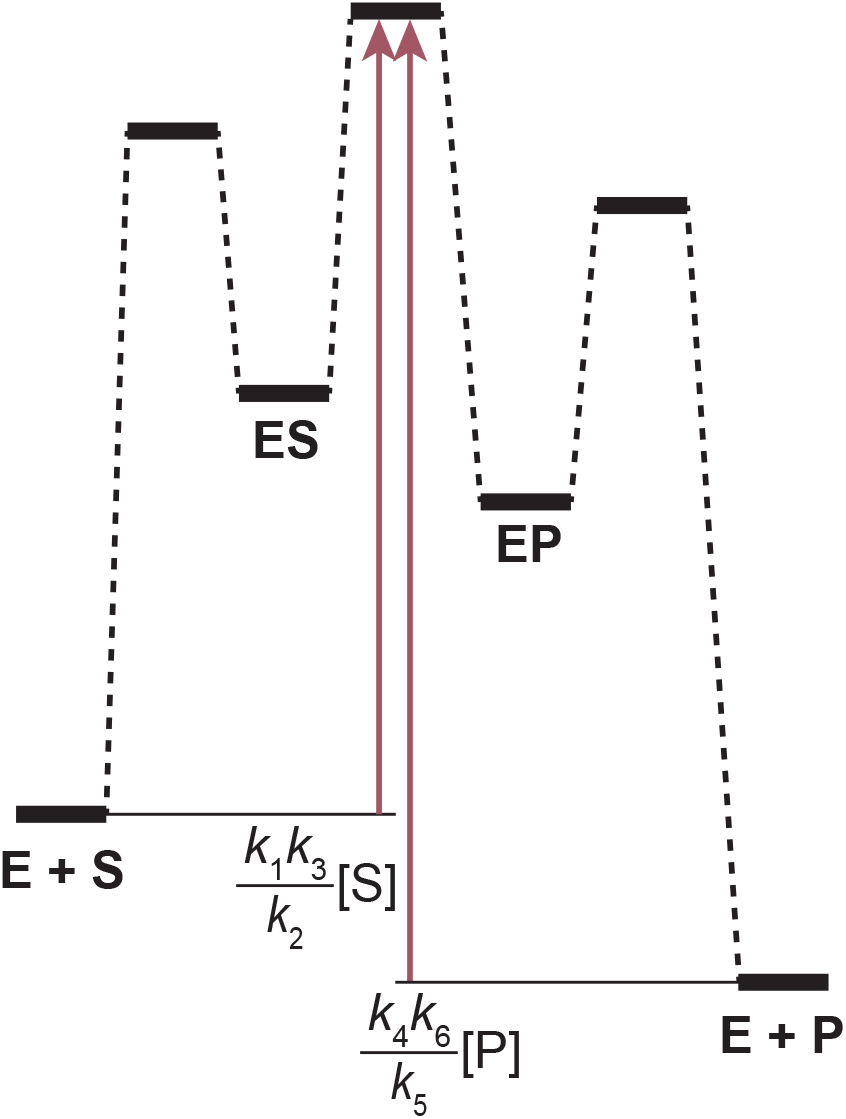
*k*_cat_/*K*_M_ of the forward reaction and the reverse reaction of a reversible reaction. The transitions state for the chemical conversion step (ES → EP) is the highest, and the height of the transition state relative to E + S and E + P are indicated by teal arrows corresponding to *k*_cat_/*K*_M_ for the forward reaction and the reverse reaction, respectively.

Now using the above two equations, one can show that the ratio of (*k*_cat_/*K*_M_)_f_ to (*k*_cat_/*K*_M_)_r_ is identical to *K*_eq_, the equilibrium constant of the reaction:

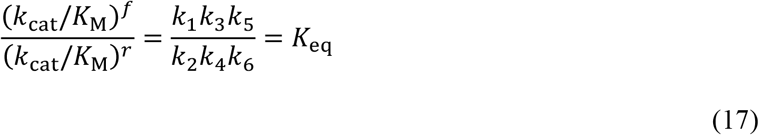

One can easily verify that this relationship holds when the highest transition state is the substrate association step (E + S ⇄ ES) or the product release step (EP ⇄ E + P). Whatever transition state is chosen, (*k*_cat_/*K*_M_)_f_ includes all the elementary rate constants from E + S to the transition state, and (*k*_cat_/*K*_M_)_r_ includes all the elementary rate constants from E + P to the same transition state. As shown in Eq. 17, the ratio of (*k*_cat_/*K*_M_)_f_ to (*k*_cat_/*K*_M_)_r_ contains the product of all the forward rate constants in the denominator and the product of all the reverse rate constants in the numerator, which is equivalent to *K*_eq_.

One can also prove the Haldane relationship by simply calculating the ratio of (*k*_cat_/*K*_M_)_f_ to (*k*_cat_/*K*_M_)_r_ in Eqs. 14 and 15 without assuming any single transition state is significantly higher than other transition states. Because the forward reaction and the reverse reaction go through the same transition states simply in the reverse order, the relative contribution of each transition state is identical to (*k*_cat_/*K*_M_)_f_ and (*k*_cat_/*K*_M_)_r_. Because *k*_cat_/*K*_M_ is expressed as the sum of the kinetic terms corresponding to the transition states (Eq. 9), reversing the order of the kinetic terms in the expression does not change the sum. The only difference between (*k*_cat_/*K*_M_)_f_ and (*k*_cat_/*K*_M_)_r_ is the starting point (or the reference level for the transition states), E + S and E + P, respectively, and this difference makes the ratio of the two parameters *K*_eq_.

As we stated already, the same step is rate-limiting for both (*k*_cat_/*K*_M_)_f_ to (*k*_cat_/*K*_M_)_r_. If the substrate-association step is rate-limiting for (*k*_cat_/*K*_M_)_f_, the same step, which is the product-release step in the reverse reaction, is also rate-limiting for (*k*_cat_/*K*_M_)_r_. From this principle, one can deduce intuitively that it is not possible for an enzyme to have *k*_cat_/*K*_M_ at the diffusion limit for both forward and reverse reactions, which we have proved above using the Haldane relationship. If (*k*_cat_/*K*_M_)_f_ is at the diffusion limit, the substrate-association step is rate-limiting for (*k*_cat_/*K*_M_)_f_, which means that the product-release step should be rate-limiting for (*k*_cat_/*K*_M_)_r_. Therefore, the substrate-association step cannot be rate-limiting for (*k*_cat_/*K*_M_)_r_.^10^

## Conclusion

*k*_cat_/*K*_M_ is frequently misunderstood as a composite kinetic parameter derived from *k*_cat_ and *K*_M_. Even many biochemistry textbooks wrongfully state that *k*_cat_/*K*_M_ is a way to consider both catalytic power (*k*_cat_) and binding affinity (*K*_M_) of the enzyme for its substrate. Different from *k*_cat_ and *K*_M_, *k*_cat_/*K*_M_ has a clear and straightforward meaning that can be visualized in the reaction energy diagram; *k*_cat_/*K*_M_ corresponds to the height of the highest transition state that the enzyme needs to cross along the reaction pathway of the enzyme-catalyzed reaction. Even in a minimal kinetic model as Model 1, *k*_cat_/*K*_M_ can be limited by the substrate association, the chemical conversion, or even the product release. Thus, using *k*_cat_/*K*_M_ to consider both catalytic power and binding affinity is misleading. For example, it is quite common that *k*_cat_/*K*_M_ is limited by the substrate association, i.e. *k*_cat_/*K*_M_ ~ *k*_1_, and *k*_1_ has nothing to do with the catalytic power or the binding affinity of the enzyme. The proper understanding of *k*_cat_/*K*_M_ using the reaction energy diagram is extremely helpful to analyze the effects of mutations in the enzyme, structural variations in the substrate, modification of the experimental condition, and so forth. When any of these changes affects the energy of the highest transition state, we observe a change in *k*_cat_/*K*_M_. And, the extent of the change we observe in *k*_cat_/*K*_M_ provides valuable information on the reaction energetics and catalytic mechanism of the enzyme. Moreover, the primary mechanism for the enzyme to catalyze a reaction is to lower the energy of the transition states, and *k*_cat_/*K*_M_ is a direct way to observe the outcome of this stabilization (15, 16). This is why enzymologists consider *k*_cat_/*K*_M_ along with *k*_cat_ as a fundamental kinetic parameter in enzyme kinetics and consider *K*_M_ as simply the ratio of the two fundamental kinetic parameters (1, 5, 6). Finally, I want to point out that the common practice to determine *k*_cat_ and *K*_M_ first by fitting the Michaelis–Menten equation to the experimental data and calculate *k*_cat_/*K*_M_ later using the two fitted parameters is not desirable. This practice produces a *k*_cat_/*K*_M_ value with an unnecessarily exaggerated fitting error due to the error propagation. Considering that *k*_cat_/*K*_M_ is such a fundamental kinetic parameter, one should avoid the error propagation by determining *k*_cat_ and *k*_cat_/*K*_M_ directly from the fitting of the Michaelis–Menten equation with the following fitting function: *f* = *a*\**x*/(*a*/*b* + *x*), where *f* = *v*/E_t_, *x* = [S], *a* = *k*_cat_ and *b* = *k*_cat_/*K*_M_. Then, one can calculate *K*_M_ from *k*_cat_ and *k*_cat_/*K*_M_, only if it is necessary.

## Acknowledgements

I originally conceived the idea of this manuscript from my discussion with Dr. Dexter B. Northrop (University of Wisconsin) on the reciprocal expression of *k*_cat_/*K*_M_ during my PhD thesis examination in 2000. I thank Dr. Andy Hudmon (Purdue University) for encouraging me to publish this manuscript and giving me critical suggestions on the initial draft. I also thank Drs. Rong Huang, Casey Krusemark, and Zhong-Yin Zhang for their thoughtful suggestions on the manuscript.

## Footnotes

1 The traditional form of the Michaelis-Menten equation was written with *V*_max_, instead of *k*_cat_E_t_, as: 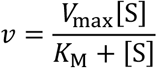 The main reason of the popular use of *V*_max_ instead of *k*_cat_ in the past was that the concentration of the total active enzyme (E_t_) was unknown or inaccurate in many cases. For the same reason, *V*/*K* was more popularly used than *k*_cat_/*K*_M_. While *k*_cat_ and *k*_cat_/*K*_M_ are kinetic parameters characteristic for an enzyme and a substrate, *V*_max_ and *V*/*K* are kinetic parameters determined with an enzyme at an arbitrary concentration. As protein purification techniques have advanced and the analytical methods for protein concentration determination have improved, *V*_max_ and *V*/*K* are often replaced with *k*_cat_ and *k*_cat_/*K*_M_.

2 Many biochemistry textbooks use an even simpler kinetic model for the derivation of the Michaelis– Menten equation: 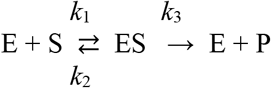 However, this kinetic model is not suitable in dissecting the meanings of *k*_cat_ and *k*_cat_/*K*_M_ rigorously because the chemical conversion step (ES ⇄ EP) and the product release step (EP → E + P) are not separated, and *k*_cat_ is determined by a single elementary rate constant (*k*_3_)! The separation of the two steps is indeed necessary because the product release, not the chemical conversion, limits the overall turnover in many enzyme-catalyzed reactions (1).

3 The rate equation can be derived either with the King–Altman method (2) or with the net rate constant method (3). Especially, the inclusion of the single irreversible step for the product release in Model 1 allows us to use the net rate constant method for the derivation of the rate equation.

4 The expression for *K*_M_ for Model 1 is even more complex than those for *k*_cat_ or *k*_cat_/*K*_M_: 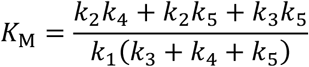

5 As the name implies, the pseudo-first-order rate constant is not a true rate constants; when [S] varies, the constant varies accordingly. In enzyme kinetics, however, we treat the pseudo-first-order rate constants as same as any other first-order rate constants because typically we deal with the kinetics of enzymatic catalysis at a fixed concentration of [S]. Treating [S] as a constant is also a common practice in experimental determination of the initial velocity (*v*_0_); the initial velocity should be determined within a timeframe in which [S] does not change significantly.

6 We can demonstrate the mathematical relationship between the heights of the transition states and *k*_1_*k*_3_/*k*_2_ and *k*_1_*k*_3_*k*_5_/*k*_2_*k*_4_, using the property that the barrier height in the reaction energy diagram is linearly proportional to ln(1/*k*). 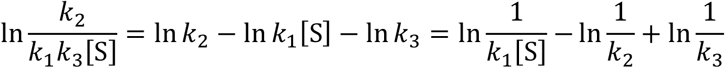 The three terms on the right side are linearly proportional to the barrier heights corresponding to *k*_1_[S], *k*_2_, and *k*_3_, respectively. By subtracting and adding the arrow lengths in the reaction energy diagram in Figure 2, one can visually verify that *k*_1_*k*_3_/*k*_2_ corresponds to the height of the transition state for the chemical conversion step (ES → EP) relative E + S. Using the same approach, we can demonstrate that *k*_1_*k*_3_*k*_5_/*k*_2_*k*_4_ corresponds to the height of the transition state for the product release step (EP → E+P) relative E + S.

7 ‘Rate-limiting’ is not a suitable word for *k*_cat_/*K*_M_ because *k*_cat_/*K*_M_ is not a rate but a rate constant. I still use ‘rate-limiting’ in this manuscript simply following the convention.

8 The rate-limiting step for *k*_cat_/*K*_M_ is not necessarily the rate-limiting step for *k*_cat_. At a saturating concentration of the substrate, the amount of the free enzyme (E) is negligible. i.e. the bound states of the enzyme (ES and EP in Model 1) are highly populated, and the highest transition state may not have the dominant influence on *k*_cat_. For example, the substrate association step cannot not affect *k*_cat_, even when its transition state is the highest. Moreover, the application of the concept of the rate-limiting step to *k*_cat_ is complicated and open to question (4).

9 Practically, it is quite unlikely to raise the energy of a trough without affecting the height of neighboring transition states because the structural similarity exists between the intermediate and the transition states. The treatment of selectively raising the energy of a trough here is only for a conceptual assessment of the property of *k*_cat_/*K*_M_.

10 One can think of an extreme case in which the transition states for the substrate association step and for the product release step have the same height, and the equilibrium constant of the reaction is 1. Even in this extreme case, *k*_cat_/*K*_M_ for both forward and reverse reactions can reach up to only half of the diffusion limit, because two transition states have the same contribution to *k*_cat_/*K*_M_ (Eq. 9 and Table 1).

